# Identification of potential colorectal-cancer-protective bioactive natural compounds in the human gut microbiome

**DOI:** 10.64898/2026.02.17.706414

**Authors:** Daniela Leuzzi, Simone Rampelli, Castrense Savoiardo, Daniel Scicchitano, Federica D’Amico, Teresa Frisan, Pierluigi Martelli, Marco Candela, Silvia Turroni

## Abstract

Increasing evidence links the gut microbiome to the development and progression of colorectal cancer (CRC). Despite the remarkable diversity of microbial biosynthetic gene clusters (BGCs), no CRC-protective bioactive natural compounds originating from gut microbiome BGCs have yet been detected. Here, we investigated BGCs within *de novo* gut metagenome-assembled species reconstructed from 559 individuals, including healthy controls, adenoma and CRC patients. Comparative analysis revealed distinct BGC patterns across groups, with four BGCs being significantly enriched in healthy individuals. One of these was predicted to be involved in the biosynthesis of a metabolite closely related to the anticancer compound curacin-A. Notably, this BGC was also overrepresented in healthy centenarians, suggesting a role in achieving long-term cancer-free health. Our findings highlight the importance of exploring gut microbiome BGCs as a source of novel bioactive molecules for cancer prevention and treatment, as well as for other health purposes.

## Introduction

Colorectal cancer (CRC) is the second deadliest cancer globally (Hossain *et al*., 2022). Within the European Union, it is the second most common cancer among women and the third most common among men (Hossain *et al*., 2022; Roshandel *et al*., 2024; Candela *et al*., 2011). In 2020 alone, 19.3 million new cases have been estimated, resulting in 10 million cancer-related deaths. In future decades, both incidence and fatalities are expected to rise (Hossain *et al*., 2022; Roshandel *et al*., 2024). It is estimated that there will be a worldwide increase of about 63.3% in the number of new cases by 2040, with slight declining trends expected in high-risk countries and opposite strong increasing trends in low-risk countries, due to socioeconomic factors (Roshandel *et al*., 2024). The risk of developing CRC is influenced by both non-modifiable and modifiable factors. The latter include cigarette smoking, a sedentary lifestyle, psychological stress, the regular and prolonged consumption of alcohol, an imbalanced diet (Aykan 2015; Roshandel *et al*., 2024), and dysbiotic changes in the gut microbiome (GM) (Gausman *et al*., 2020). Indeed, GM compositional alterations have emerged as a possible driver of CRC onset (Candela *et al*., 2011; Masheghati *et al*., 2024; Kim & Lee, 2022). Studies have consistently identified CRC-promoting taxa, such as *Escherichia coli, Bacteroides fragilis, Fusobacterium nucleatum*, enterotoxigenic *Bacteroides fragilis, Helicobacter pylori, Salmonella* and *Enterococcus faecalis* (Pandey *et al*., 2024; Candela *et al*., 2011). The underlying mechanisms include the induction of chronic inflammation, the modulation of immune responses and, not least, the promotion of genotoxicity. For instance, some microbes can produce toxins that damage DNA (e.g. *E. coli* and *Salmonella*) (Gradel *et al*., 2009; Escobar-Páramo *et al*., 2004) or activate inflammatory (e.g. *B. fragilis*) and immune signals (e.g. *F. nucleatum*), creating a microenvironment favorable to tumor initiation and progression (Sobhani *et al*., 2011; Morgillo *et al*., 2018). Conversely, it is known that certain commensal taxa - such as *Faecalibacterium, Eubacterium, Bifidobacterium*, and *Lactobacillus* - have protective effects against CRC (Candela *et al*., 2011; Pandey *et al*., 2024). These effects are likely to be mediated by the competitive exclusion of CRC-promoting taxa, the reinforcement of the intestinal barrier, and the maintenance of a homeostatic immune environment. The responsible molecules include short-chain fatty acids (SCFAs), primarily acetate, propionate, and butyrate, which are generated through the GM fermentation of dietary fiber (Louis & Flint, 2009). Notably, butyrate has been shown to exhibit potent anti-inflammatory and anti-proliferative properties, positioning it as a key GM metabolite in CRC prevention (Chen & Li, 2020). In addition to SCFAs, indoles, resulting from tryptophan metabolism, have anti-cancer and anti-inflammatory effects against CRC (Liu *et al*., 2023). However, to the best of our knowledge, no bioactive natural compounds (BNCs), defined as microbially produced chemicals affecting other organisms or biological systems, with CRC-protective functions have yet been identified within the human GM.

In bacteria and fungi, the biosynthesis of BNCs is governed by mega-clusters of co-localized genes known as biosynthetic gene clusters (BGCs). BGCs include genes that code for core multifunctional enzymes (building the scaffold), tailoring enzymes (decorating the scaffold), transporters, transcription factors and resistance mechanisms (Medema *et al*., 2015; Martinet *et al*., 2019; Mao *et al*., 2018). In particular, core genes show more aligned modules, each divided into different domains, as catalytic subunits in charge of individual biosynthetic steps, as well as other domains involved in precursor loading, condensation, modification and release (Hwang *et al*., 2020). BGCs comprise different classes, such as non-ribosomal peptides (NRPs), polyketide synthases (PKSs), hybrid PKS-NRP systems, and ribosomally synthesized and post-translationally modified peptides (RiPPs) (Dat *et al*., 2022). The remarkable microbial diversity of BGCs underlies the biosynthesis of a vast array of BNCs, each performing distinct biological functions that are potentially relevant to human health. These functions include, but are not limited to, antimicrobial, anti-inflammatory, and anticancer properties (Campos-Magaña *et al*., 2025). For instance, NRPSs and PKSs are responsible for the production of antibiotics (vancomycin (van Groesen *et al*., 2021) and erythromycin (Khabthani *et al*., 2021)) as well as immunosuppressants (cyclosporin and rapamycin) (Pogorevc & Müller, 2022). Several microbial BGCs with potent anticancer activities have also been identified (Rui *et al*., 2025), including those responsible for the biosynthesis of doxorubicin (PKS), trabectedin (alkaloid), and dolastatin (NRP) (Nakamura *et al*., 2016; Gao *et al*., 2021). Anticancer BGCs are known to act through diverse anti-proliferative mechanisms, such as: (i) nucleoside analogues that interfere with DNA synthesis; (ii) DNA-alkylating agents that disrupt DNA replication; and (iii) microtubule-binding agents that block mitosis (Chang *et al*., 2024; Dubois & Cohen, 2009; Čermák *et al*., 2020).

Members of the human microbiome, particularly the oral microbiome, have been reported to provide for more than 3,000 BGCs, many of which remain uncharacterized (Donia *et al*., 2014; Youngblut, *et al*., 2020; Aleti *et al*., 2019). For instance, genera such as *Bacteroides, Corynebacterium, Ruminococcus, Rothia, Parabacteroides, Escherichia, Lactobacillus, Enterococcus*, and *Haemophilus* possess between 2 and 7 BGCs per genome (Donia *et al*., 2014). Regarding the human GM, the vast majority of characterized BGCs have been associated with antimicrobial functions, such as lantibiotics, thiopeptides, bacteriocins, microcins, and thiazole/oxazole-modified microcins (TOMMs) (Donia & Fischbach, 2015). In a recent large-scale meta-analysis of the biosynthetic potential of RiPPs in human-associated bacteria, Zhang *et al*. (2025) uncovered 30 actively transcribed RiPP families that define the healthy GM. These families showed consistent negative associations with multiple diseases, including CRC. Although these landmark discoveries underscore the pivotal role of microbiome-derived BGCs in human health, CRC-protective BNCs originating from GM BGCs have not yet been identified.

Given that anticancer activities are common among microbial BGCs, we hypothesize that the human GM may serve as an untapped endogenous sink of BGCs providing BNCs that could protect against cancer, including CRC. Here, we conducted a targeted bioinformatic meta-analysis of studies examining GM changes associated with CRC onset (Zeller *et al*., 2014; Feng *et al*., 2015; Vogtmann *et al*., 2016; Yu *et al*., 2017; Hannigan *et al*., 2018) with the aim to identify BGCs enriched in the GM of healthy individuals as compared to individuals with CRC or colorectal adenomas. Using this approach, we successfully identified BGCs associated with a healthy GM that encode putative anti-carcinogenic compounds. Our findings suggest the existence of a new, microbiome-mediated defence system against CRC onset.

## Results

### Identification of BGCs characterizing the gut metagenome of healthy individuals compared to CRC and adenoma populations

A total of 16,936 metagenome-assembled genomes (MAGs), reconstructed from 559 human gut metagenomes (including 258 from individuals with CRC, 81 from individuals with colorectal adenomas, and 220 from healthy individuals) from 5 publicly available, geographically diverse metagenomic studies of CRC (Zeller *et al*., 2014; Feng *et al*., 2015; Vogtmann *et al*., 2016; Yu *et al*., 2017; Hannigan *et al*., 2018), were analyzed to predict and annotate BGCs. A total of 29,238 BGCs were detected, the majority of which were classified as RiPPs (n=23,629), followed by NRPSs (n=2,284), terpenes (n=657) and PKSs (n=196). BGCs were clustered into 1,148 BGC families based on a sequence similarity threshold of 70%. BGC families showing more than twofold enrichment in MAGs from healthy individuals compared to MAGs from CRC/adenoma patients were considered characteristic of a healthy GM (and potentially protective against CRC) and were selected for identification. This led to 118 BGC families being identified, 5 of which were exclusively detected in MAGs from healthy individuals. Three of these 5 families were unclassified, while 2 were assigned to the NRPS and RiPP categories, respectively. The core genes from the 118 BGC families were retrieved, and their respective abundances in the 559 human gut metagenomes were quantified (**Figure 1**). Four core genes showed significantly higher abundances in gut metagenomes from healthy individuals compared to CRC/adenoma patients (p<0.05, Wilcoxon rank sum test). The corresponding BGCs, defined as healthy microbiome (HM)-BGCs, were assigned to the following classes: RiPP-like (HM-BGC1), RiPP Recognizing Element (RRE) (HM-BGC2), NRP (HM-BGC3) and β-lactone (HM-BGC4) (**Figure 2**). The MAGs carrying the 4 HM-BGCs were retrieved and identified taxonomically (**Table 1**). The following species were identified: *Streptococcus salivarius* (HM-BGC1), *Blautia obeum* (HM-BGC2), *Eubacterium hallii* (HM-BGC3) and *Eubacterium eligens* (HM-BGC4). Interestingly, when assessing the prevalence and abundance of these taxa across the 559 human gut metagenomes, we found no significant differences between the study groups (p>0.05). This suggests that the potential CRC-protective effects may depend on the specific BGC cargo carried by GM taxa, rather than on the taxonomic composition alone.

**Table 1.** Main features of healthy microbiome-biosynthetic gene clusters (HM-BGCs). For each of the 4 HM-BGC identified in the present study, the respective taxon, prevalence (expressed as a percentage), mean (expressed as reads per kilobase of gene per million mapped reads, RPKM), and standard deviation in gut metagenomes of healthy individuals and individuals with colorectal cancer (CRC) and adenomas, are reported.

| BGC_id | Taxon | Prevalence |  |  | Mean (RPKM) |  |  | Standard deviation (RPKM) |  |  |
| --- | --- | --- | --- | --- | --- | --- | --- | --- | --- | --- |
|  |  | % CRC | % adenoma | % healthy | CRC | adenoma | healthy | CRC | adenoma | healthy |
| HM-BGC1 | *Streptococcus salivarius* | 98.83 | 100 | 99.55 | 23.62 | 46.941 | 33.85 | 66.67 | 141.64 | 79.40 |
| HM-BGC2 | *Blautia obeum* | 99.61 | 100 | 100 | 69.22 | 215.69 | 123.14 | 130.01 | 243.89 | 204.84 |
| HM-BGC3 | *Eubacterium hallii* | 98.06 | 98.76 | 99.09 | 38.43 | 86.05 | 61.61 | 74.28 | 107.8 | 91.64 |
| HM-BGC4 | *Eubacterium eligens* | 96.12 | 95.06 | 96.82 | 54.94 | 30.28 | 76.27 | 111.53 | 81.4 | 135.34 |

**Figure 1.**
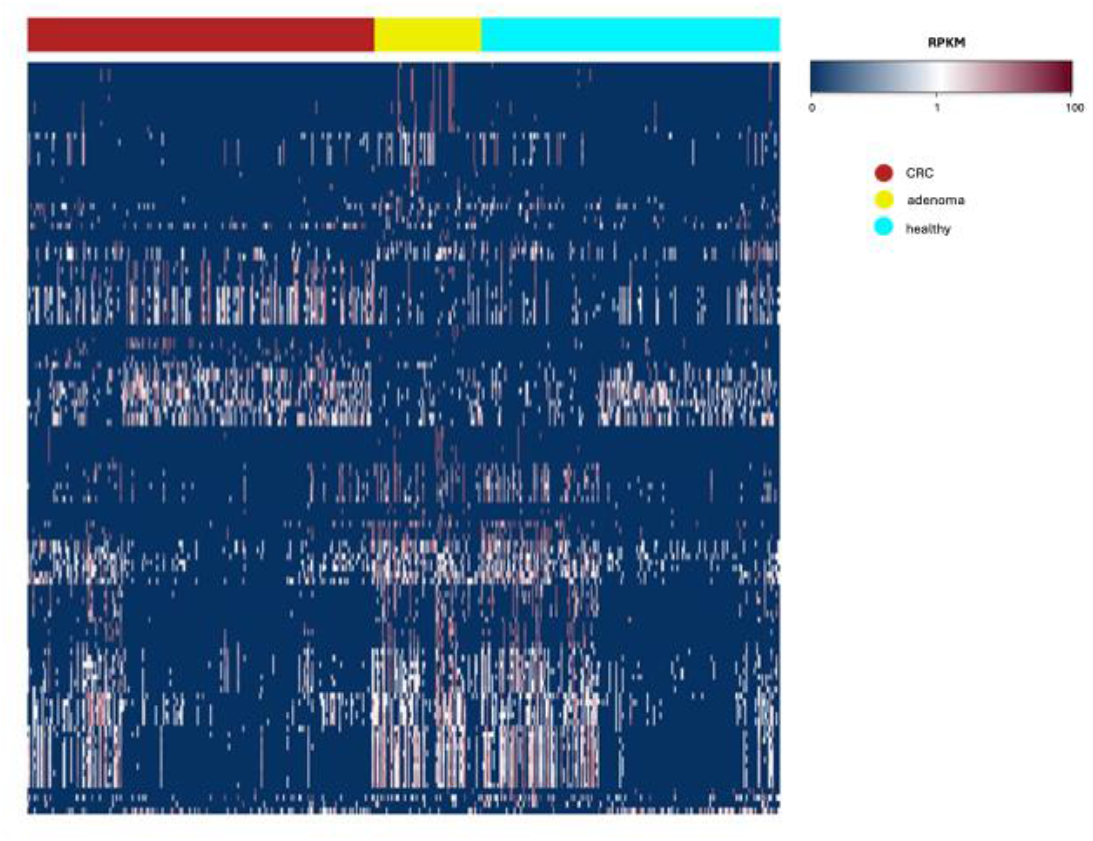
Abundance distribution of the core genes of the biosynthetic gene cluster families identified in the gut metagenomes of healthy individuals and individuals with colorectal cancer or adenomas. Heatmap showing the abundance (expressed in reads per kilobase of gene per million mapped reads, RPKM) of the core genes of the 118 BGC families identified in the gut metagenomes of healthy individuals and individuals with colorectal cancer (CRC) or adenomas.

**Figure 2.**
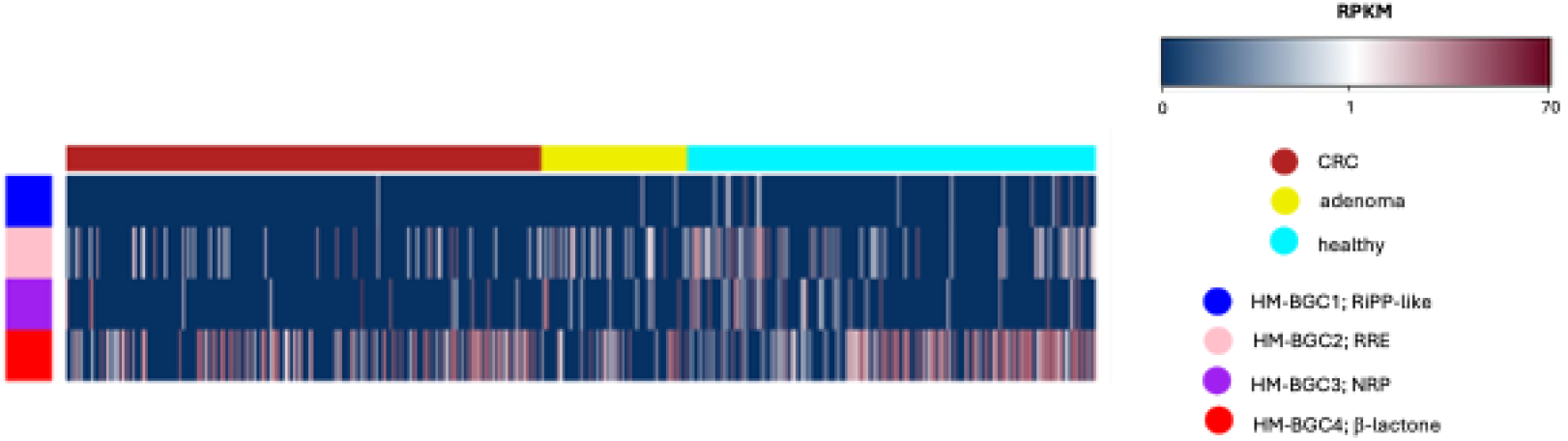
Abundance distribution of the biosynthetic gene clusters identified as more prevalent in the gut metagenomes of healthy individuals compared to individuals with colorectal cancer or adenomas. Heatmap showing the abundance (expressed in reads per kilobase of gene per million mapped reads, RPKM) of the 4 HM-BGCs that were significantly more abundant in healthy individuals compared to individuals with colorectal cancer (CRC) or adenoma (p<0.05, Wilcoxon rank sum test). For each HM-BGC, the corresponding class is reported: ribosomally synthesized and post-translationally modified peptide (RiPP)-like (HM-BGC1), RiPP Recognizing Element (RRE) (HM-BGC2), non-ribosomal peptide (NRP) (HM-BGC3) and β-lactone (HM-BGC4).

### Synteny and putative function of the Healthy Microbiome-BGCs

For each HM-BGC, the gene organization and composition were analyzed and, when possible, the corresponding BNC was predicted (**Figure 3**). HM-BGC1 is a RiPP-like cluster containing 2 bacteriocin IIc homologs. HM-BGC2, classified as RRE-like, displays an arrangement consisting of an ABC transporter followed by a radical S-adenosyl-L-methionine (rSAM)enzyme. HM-BGC3 exhibits an NRPS organization, comprising a series of core biosynthetic domains (AMP-binding, condensation) accompanied by tailoring enzymes such as methyltransferase, phosphopantetheinyl transferase and thioesterases, as well as genes associated with cyclic-lactone autoinducers (AgrB, TIGR04223). Finally, HM-BGC4, a β-lactone cluster, exhibits an extended architecture featuring regulatory elements (AraC family transcriptional regulator), an additional biosynthetic gene (alkyl hydroperoxide reductase subunit), and 2 core biosynthetic genes (HMGL-like AMP-binding proteins), together with additional cyclic-lactone autoinducer-associated genes (AgrB, TIGR04223). The synteny of the four HM-BGCs was compared with that of their closest homologs in the BGC Atlas database. HM-BGC1 was assigned to family GCF_352, HM-BGC2 to GCF_2325, HM-BGC3 to GCF_14140, and HM-BGC4 to GCF_9548. The comparative analysis confirmed that HM-BGC1 is a RiPP-like cluster that encodes the bacteriocin IIc of *S. salivarius*, which is already represented in existing databases. HM-BGC2 corresponded to an incomplete region within known sactipeptide-producing BGCs from members of the *Lachnospiraceae* family (to which *B. obeum* belongs). HM-BGC3 shared both sequence homology and synteny with NRPS-type BGCs belonging to the BGC Atlas family GCF_14140, which are mostly assigned to species of the *Lachnospiraceae* family (including *E. hallii*). Finally, HM-BGC4 belonged to a relatively heterogeneous family composed entirely of BGCs from *E. eligens*.

**Figure 3.**
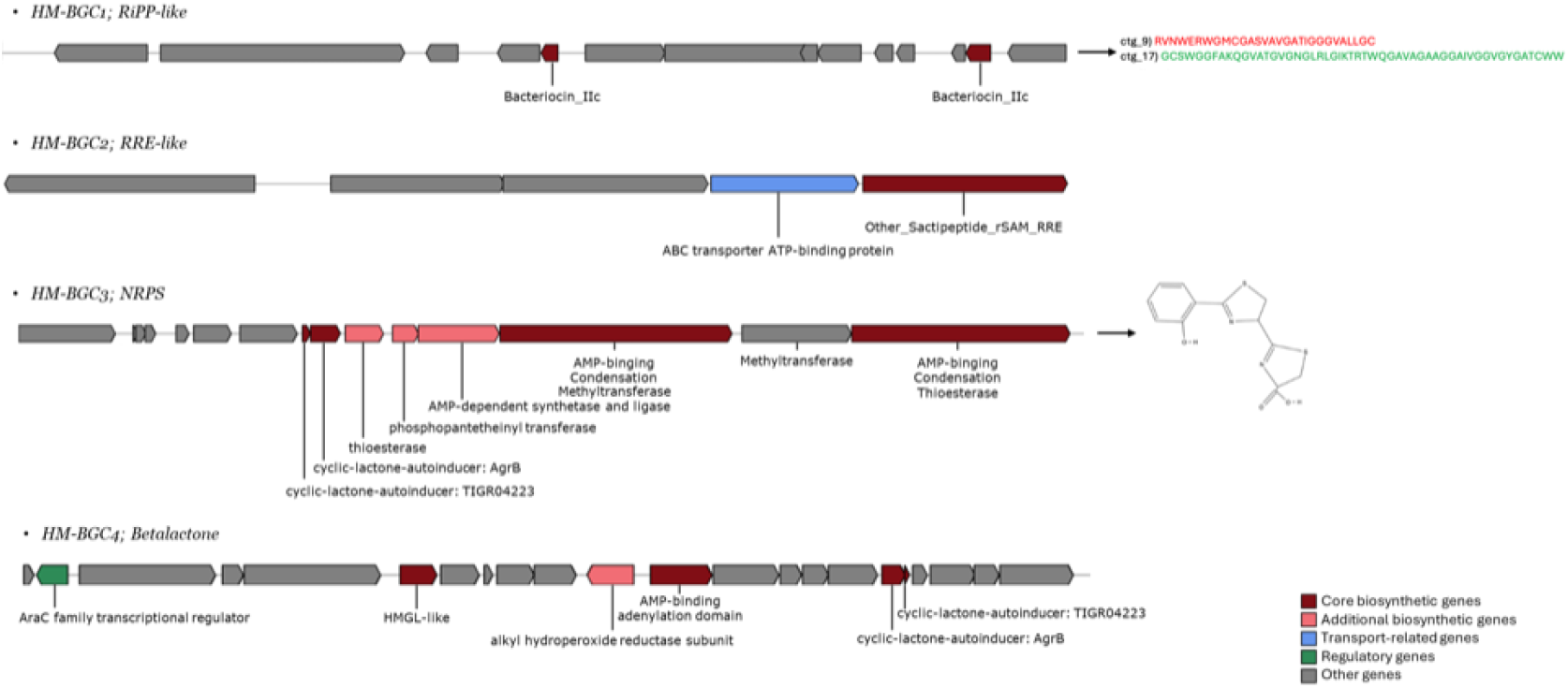
Synteny of the 4 healthy microbiome-biosynthetic gene clusters (HM-BGCs). Genes are colored according to their predicted functions: core biosynthetic genes (dark red), additional biosynthetic genes (salmon), transport-related genes (blue), regulatory genes (green), and other genes (gray). For each HM-BGC, the corresponding class is reported: ribosomally synthesized and post-translationally modified peptide (RiPP)-like (HM-BGC1, signal peptides of the two bacteriocin IIc are highlighted in red and green), RiPP Recognizing Element (RRE) (HM-BGC2), non-ribosomal peptides (NRP) (HM-BGC3) and β-lactone (HM-BGC4). The chemical structure shown corresponds to the putative bioactive natural compound predicted for HM-BGC3.

Where possible, the chemical structures of the encoded metabolites were inferred using PRISM (Skinnider *et al*., 2017) and RIPPMiner (Agrawal *et al*., 2017). A putative BNC could only be identified for HM-BGC3. This consists of a bicyclic heterocyclic core of 2 thiazoline rings, ones carrying a carboxylic acid group, forming a bicyclic system connected to a phenyl ring bearing a phenolic hydroxyl group. Overall, the structure combines acidic functionalities (carboxyl and phenolic hydroxyl) with a sulfur- and nitrogen-rich heterocyclic scaffold, giving it a mixed aromatic– heteroaromatic character with potential biological activity. Interestingly, the core gene of HM-BGC3 showed 56% sequence similarity and 36% sequence identity to *CurF* (**Supplementary File 1**). CurF is a key enzyme of curacin A BGC from *Moorea producens* (Chang *et al*., 2004) and is responsible for forming the characteristic curacin thiazoline ring. This homology suggests that the HM-BGC3 core gene may encode a curacin-related thiazoline-forming enzymatic machinery; we refer to this henceforth as the “*CurF-like*” HM-BGC3 core gene.

### Possible evolutionary origin of the “*CurF-like*” HM-BGC3 core gene

Based on the sequence of the “*CurF-like*” HM-BGC3 core gene, we first delineated the modular organization of the encoded “CurF-like” protein. We identified a canonical NRPS/PKS architecture comprising an N-terminal condensation domain, an AMP-binding adenylation domain and a C-terminal S-adenosylmethionine-dependent methyltransferase (MT) domain (**Figure 4A**). To gain structural insight into this multidomain enzyme, we generated an *in-silico* model using AlphaFold (AF) (Jumper *et al*., 2021). The resulting structure of the “CurF-like” protein was compared with homologous modules from curacin A BGCs. We assessed the superimposition in terms of TM-score, ranging between 0 and 1 (where 1 indicates a perfect match between two structures), and Root Mean Square Deviation (RMSD). Structural alignment revealed a high degree of similarity to the corresponding region of CurF from the cyanobacterial taxon *Lyngbya majuscula* (also known as *M. producens*) (residues 1990–3154; UniProt Q6DNE7; TM-score 0.90; RMSD 3.74 Å) (**Figure 4B**) and to the CurJ adenylation domain from *M. producens* (PDB 5THZ_A; UniProt F4Y426; residues 1269–1649; TM-score 0.88; RMSD 3.02 Å) (**Figure 4C**). These comparisons indicate that the “*CurF-like*” HM-BGC3 core gene exhibits a conserved NRPS/PKS fold, which is closely related to curacin assembly line enzymes. This supports the hypothesis that it is involved in the biosynthetic pathway of a structural analogue of curacin A (Chang et al., 2004).

**Figure 4.**
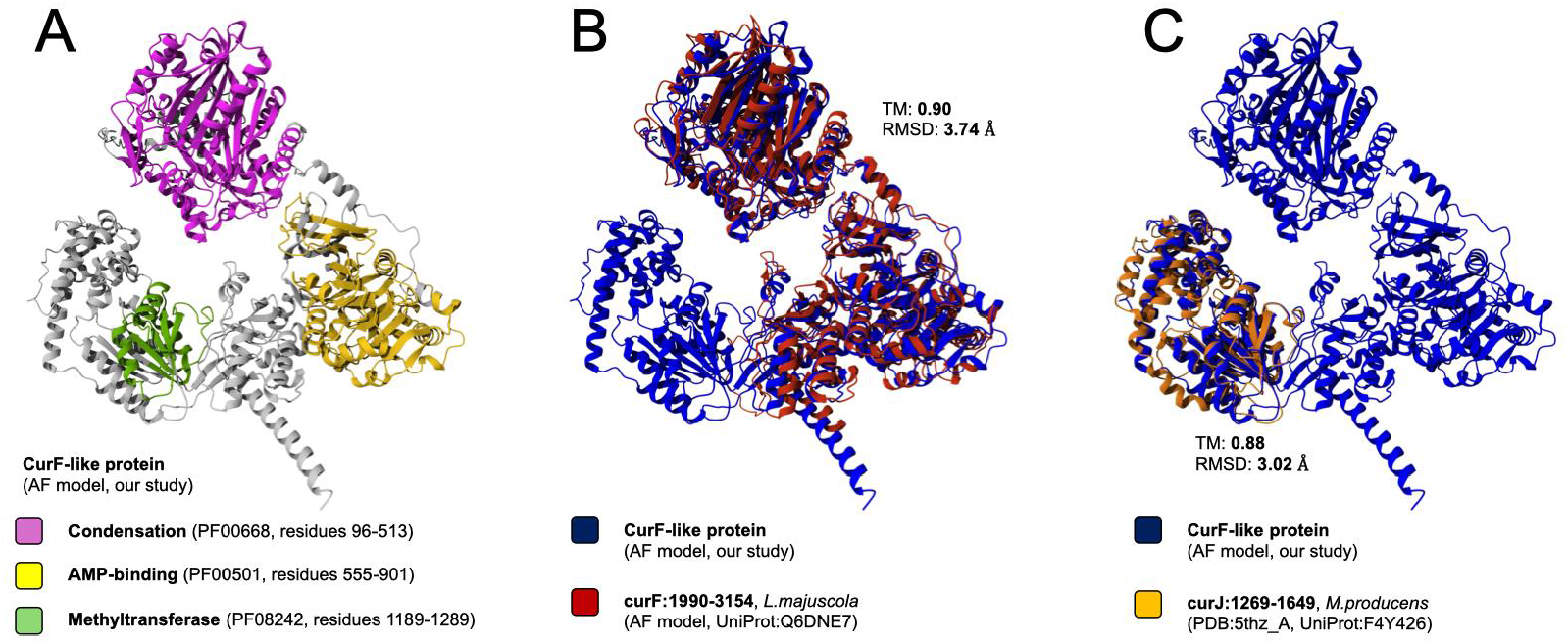
Structural model of the “CurF-like” protein and comparison with the CurF and CurJ homologous modules. **A**, Structural model of the “CurF-like” protein generated using AlphaFold2. The three predicted functional domains mapped onto the 3D structure are highlighted: the condensation domain (PF00668, residues 96-513), the AMP-binding domain (PF00501, residues 555-901), and the methyltransferase domain (PF08242, residues 1189-1289). Structural superimposition of the “CurF-like” protein model and the AlphaFold-generated model of CurF from *Lyngbya majuscula* (UniProt: Q6DNE7; residues 1990-3154), (**B**) and the experimental structure of the adenylation domain of CurJ from *Moorea producens* (UniProt: F4Y426; PDB: 5THZ, chain A; residues 1269-1649) (**C**). For each superimposition, the TM-score and Root Mean Square Deviation (RMSD) are reported.

To elucidate the evolutionary origin of the “*CurF-like*” core gene in HM-BGC3, we systematically retrieved all homologous sequences belonging to the BGC Atlas family GCF_14140 and inferred their relationships through phylogenetic reconstruction. The resulting tree incorporated the well-characterized *CurF* gene from *M. producens* as a reference (**Figure 5**), enabling a comparative framework for assessing the diversification of this enzyme family. Our phylogenetic analysis revealed that the “*CurF-like*” HM-BGC3 core gene clustered with homologous genes from *Lachnospiraceae* family members, which typically dominate the human adult GM. These include *Agathobacter rectalis, Anaerobutyricum hallii* (formerly *E. hallii*), multiple species of *Blautia*, and *Mediterraneibacter*. Notably, sequences homologous to *CurF* from *M. producens* were exclusively detected in host-associated bacterial taxa of predominantly gastrointestinal origin. This phylogenetic incongruence suggests that the “*CurF-like*” biosynthetic capacities may have originated through a horizontal gene transfer (HGT) event from cyanobacteria (specifically *M. producens*) to gut-associated bacteria, followed by domestication and diversification into “*CurF-like”* BGCs within the GM. Supporting this hypothesis, the GC content of the “*CurF-like*” HM-BGC3 core gene was 45%, closely matching that of *M. producens* (44%) while differing from that of *E. hallii* (38%), the species to which HM-BGC3 was assigned.

**Figure 5.**
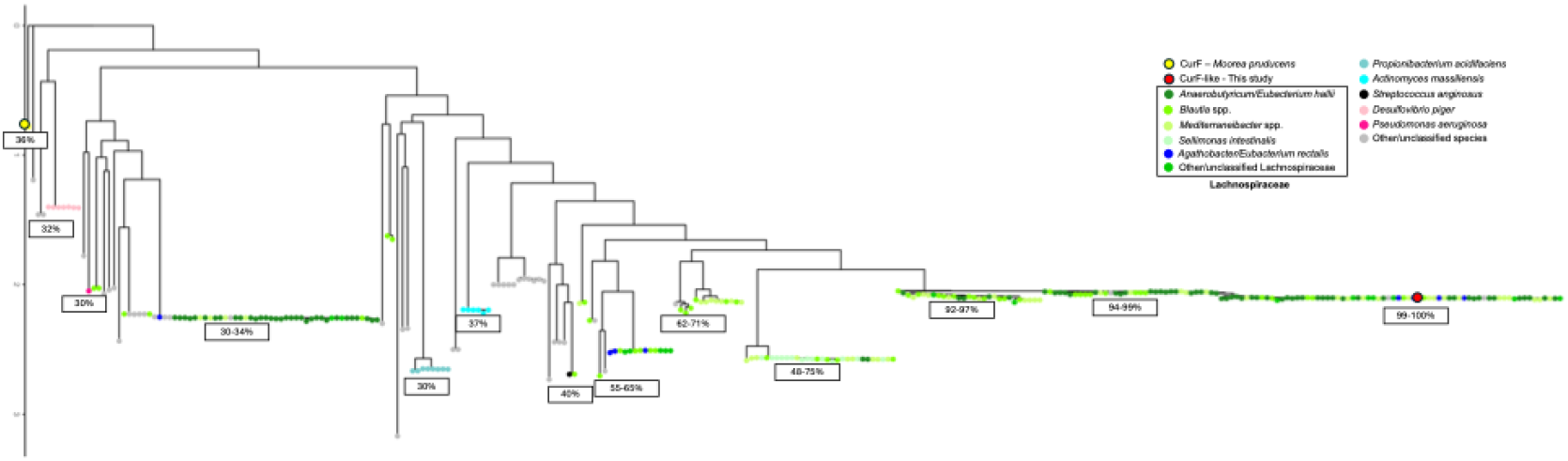
Phylogenetic tree of CurF, “CurF-like” protein from HM-BGC3 and “CurF-like” proteins from the BGC Atlas family GCF_14140. The tree shows the phylogenetic relationships between the “CurF-like” protein of HM-BGC3 (from *Eubacterium hallii*, in red), the CurF protein from the *Moorea producens* curacin A BGC (in yellow), and homologous proteins retrieved from the BGC Atlas (GCF_14140) dataset. Terminal nodes are colored according to the taxonomic identification of their source organisms, as detailed in the figure legend. Percentages next to the tips indicate the sequence identity to the “CurF-like” protein of HM-BGC3.

### Age-related distribution of the “*CurF-like*” HM-BGC3 core gene in the gut microbiome of healthy individuals

Finally, we investigated the distribution of the “*CurF-like*” HM-BGC3 core gene in the gut metagenomes of healthy individuals of different ages, namely young adults, elderly and centenarians from our previous study (Rampelli *et al*., 2020). This gene was selected for further analysis due to its relevant homology to the *CurF* and *CurJ* modules of curacin A BGCs, which are responsible for the production of curacin, a well-known anti-carcinogenic bacterial BNC (Verdier-Pinard *et al*., 1998). The choice of age groups was based on varying CRC risk: centenarians are considered a model of healthy, cancer-free aging (Marcos-Pérez *et al*., 2021), while elderly individuals and young adults are at high and low risk of developing CRC, respectively (O’Donnell, Hubbard, & Jin, 2024). Interestingly, reads mapping to the “*CurF-like*” HM-BGC3 core gene were more abundant in the gut metagenomes of both young adults and centenarians than in those of elderly individuals (mean reads per kilobase of transcript per million mapped reads (RPKM) in young adults vs. elderly vs. centenarians, 1.29 vs. 0.84 vs. 1.1) (**Figure 6**). However, this difference was not statistically significant (p>0.05, Kruskal–Wallis test) and warrants confirmation in larger cohorts.

**Figure 6.**
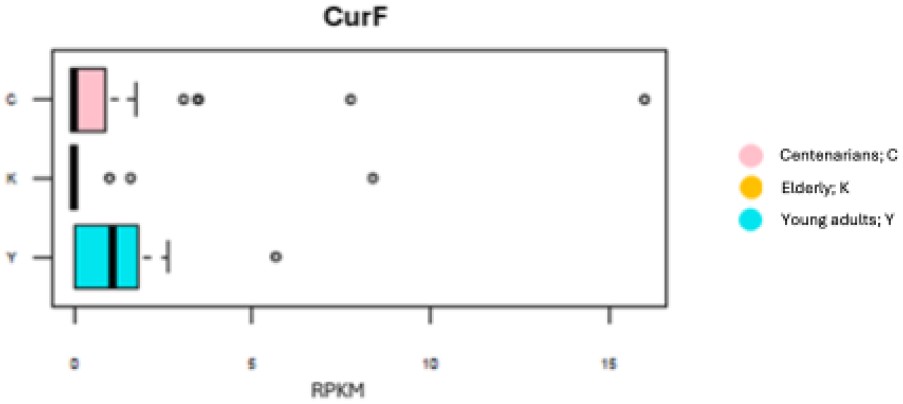
Distribution of the “*CurF-like”* core gene in the gut metagenomes of healthy individuals, including centenarians, elderly and young adults. Box plots showing the abundance distribution (expressed in reads per kilobase of gene per million mapped reads, RPKM) of the “*CurF-like*” HM-BGC3 core gene in the gut metagenomes of healthy individuals divided into the following age groups: young adults (Y), elderly (K), and centenarians (C) (p>0.05, Kruskal–Wallis test).

## Discussion and Conclusions

Through targeted bioinformatic meta-analysis of MAGs assembled from the GM of healthy individuals and individuals with CRC or colorectal adenomas (Zeller *et al*., 2014; Feng *et al*., 2015; Vogtmann *et al*., 2016; Yu *et al*., 2017; Hannigan *et al*., 2018), we identified 4 BGCs that are characteristic of healthy GM and potentially protective against CRC. These BGCs were assigned to *S. salivarius* (HM-BGC1) and to common members of the human GM, including *B. obeum* (HM-BGC2), *E. hallii* (HM-BGC3), and *E. eligens* (HM-BGC4). The HM-BGCs were classified as RiPP-like (HM-BGC1), RiPP Recognizing Element (RRE; HM-BGC2), NRP (HM-BGC3), and β-lactone (HM-BGC4), and their corresponding synteny was reconstructed. We were only able to successfully predict the corresponding BNC for HM-BGC3. Specifically, HM-BGC3 harbours a biosynthetic core gene that shows strong homology to the *CurF* and *CurJ* modules of curacin A BGC from the cyanobacterial species, *M. producens*, thus supporting its designation as a “*CurF-like*” core gene. As expected, the structure of the corresponding “CurF-like” protein displayed a high degree of similarity to the homologous regions of CurF and CurJ from curacin A BGCs, which are responsible for forming the characteristic thiazole ring of the curacin molecule. Overall, our data suggest that HM-BGC3 carries a core gene encoding a “CurF-like” protein, which is likely to be responsible for the biosynthesis of a structural analogue of curacin produced by *M. producens*. Curacin is an anticancer BNC that triggers apoptosis by inhibiting tubulin polymerization (Verdier-Pinard *et al*., 1998). No direct evidence relating to CRC is available. However, curacin A has been shown to strongly inhibit the proliferation of non-small cell lung cancer cells (Catassi *et al*., 2006), suggesting potential effects on other cancer cells.

*M. producens* is a cosmopolitan, pantropical marine cyanobacterium that is highly abundant in marine benthic environments (Engene et al., 2012). It is renowned for its remarkable biosynthetic capacity, encoding over 40 distinct BGCs, including the curacin A BGC (Ferrinho et al., 2024). Taking into account (i) the high GC content of the *“CurF-like”* gene found in gut *Lachnospiraceae* members, (ii) its phylogenetic incongruence due to its high similarity with cyanobacterial *CurF* and *CurJ*, and (iii) the notion that BGCs evolve through HGT and recombination within their modular assembly lines (Calteau *et al*., 2014), we hypothesize that HM-BGC3 in gut *Lachnospiraceae* — including the human-associated *E. hallii* — may have originated from the acquisition and subsequent domestication of the *CurF* and *CurJ* genes from the *M. producens* curacin A BGC (Calteau et al., 2014; Sélem-Mojica et al., 2019) via an ancient HGT event. Such a process could have resulted in a novel biosynthetic pathway capable of producing curacin-like structural analogues within the human — and potentially mammalian — intestinal tract.

Cyanobacteria emerged approximately 3 billion years ago and are among the most ancient and influential lineages to evolve on Earth (Philmus et al., 2025; Bhaya, Birzu, & Rocha, 2025). They are well known for contributing to numerous HGT events across phyla and even domains (Zhaxybayeva et al., 2006). Furthermore, this is not the first time that a possible HGT event between marine bacteria and human GM components has been hypothesized. Indeed, Hehemann *et al*. (2010) demonstrated that the human gut Carbohydrate Active Enzyme repertoire can be reshaped through HGT from marine bacteria, facilitated by the consumption of marine foods carrying these taxa. Similarly, an HGT event between *M. producens* and an intestinal *Lachnospiraceae* lineage could have been promoted by the ingestion of edible marine organisms that feed directly or indirectly on cyanobacteria, such as fish, mussels, or oysters (Jonasson et al., 2010). Future studies could verify this hypothesis.

Notably, when we considered gut metagenomes from an independent healthy cohort, we found that “*CurF-like*” HM-BGC3 core gene exhibited an age-related distribution, with higher levels observed in young adults and centenarians, and a marked reduction in elderlies. While it is impossible to establish prospective associations, our findings further suggest that “*CurF-like*” HM-BGC3 may play a role in promoting long-term health.

The main strength of our study is that we retrieved a large number of MAGs from 5 geographically diverse gut metagenomes, ensuring that our data are robust and independent of the so-called geographical effect (He *et al*., 2018). However, the following limitations must be acknowledged: i) some sequences were unavailable due to corruption or incompleteness; ii) some BGCs may have been missed due to the inability to reconstruct certain MAGs; iii) the sample size of the age groups in the independent healthy cohort was small, which limited statistical power; and iv) there was a lack of other host metadata with which to establish potential correlations.

In conclusion, our study led to the identification of BGCs - specifically “*CurF-like*” HM-BGC3 - that may associate with protection against CRC in human GM species. These species included typical symbionts that, however, did not differ in abundance between study groups, suggesting that the protective effect against CRC may stem from the specific combination of microbial species identity and their CRC-protective BGC repertoire. Although the results of our GM meta-analysis may open new perspectives on understanding the role of the GM in CRC, these findings remain preliminary and require validation at least at the transcriptomic and metabolic levels. Furthermore, while the rich, untapped biosynthetic potential of GM is increasingly being acknowledged (Gao *et al*., 2025), there is still a long way to go before GM-derived BNCs can be integrated into clinical applications. Specifically, future research should focus on experimentally validating predicted BNCs through BGC heterologous expression strategies (Liu *et al*., 2025), and isolation and characterization of compounds, including their bioactivity and mode of action. Once confirmed, such BNCs could be used as postbiotics in anticancer preventive and therapeutic strategies, as well as for other health-related purposes.

### Experimental Section

MAGs and raw reads were retrieved from 5 publicly available, geographically diverse metagenomic studies of CRC: Zeller *et al*., (2014), Feng *et al*., (2015), Vogtmann *et al*., (2016), Yu *et al*., (2017), and Hannigan *et al*., (2018). The populations considered in this study ranged in age from 18 to 62 years; the main characteristics of the study populations, in terms of gender, healthy vs. diseased status (CRC and adenoma), and geographical provenance are reported in **Table 2**.

**Table 2.**
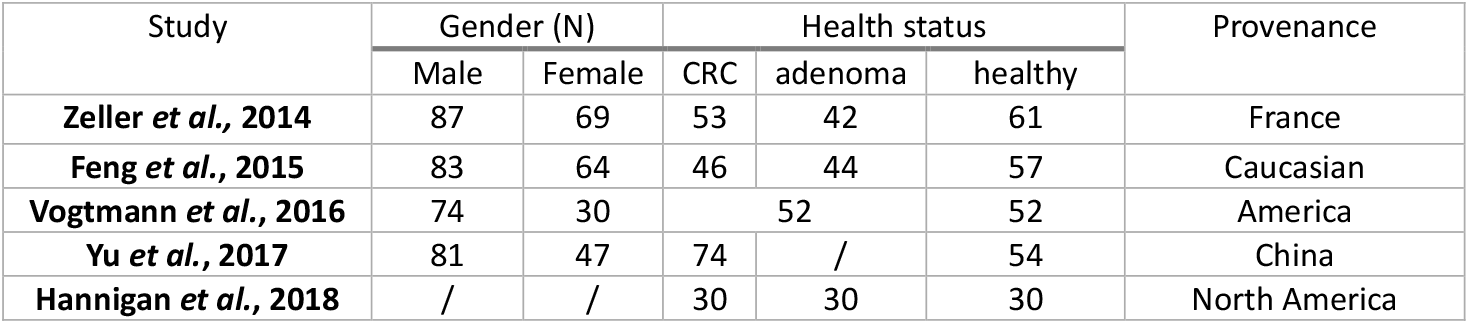
Size and characteristics of the study populations. For each population, demographic characteristics (gender, and provenance) and the number of cases (colorectal cancer – CRC and adenoma) and controls (healthy individuals) are reported.

For the analysis it was possible to retrieve a total of 559 samples (258 CRC, 81 adenomas, and 220 controls) and 16,936 MAGs classified as either high-quality (>90% complete, <5% contamination) or medium-quality (>50% complete, <5% contamination) (**Supplementary Table 1**). AntiSMASH (v. 6.1.1) (Blin *et al*., 2021) was used to identify BGCs in the MAG population. The resulting BGCs were then clustered into BGC families using BiG-SCAPE (v. 1.1.5) (Navarro-Muñoz *et al*., 2020). Family classification (‘FAM’) was retrieved, and only the GenBank (gbk) files corresponding to the clustering results were used. BGC families present exclusively or predominantly in MAGs from healthy individuals (with a prevalence of >0.70 in healthy individuals and <0.30 in individuals with CRC and adenomas) were selected. The core protocluster nucleotide sequences of these BGCs were extracted from the gbk files using AntiSMASH (v. 6.1.1, --cb-general –cb-knownclusters –cb-subclusters –cc-mibig to have a full annotation) (Blin *et al*., 2021) with the gbk2tsv tool, and a gene catalog containing the core gene sequences was created. Bowtie2 (v. 2.3.4.3, with the option ‘-- end-to-end’ ‘--very-sensitive’) (Langmead & Salzberg, 2012) was used to align the metagenome reads of each sample against the constructed proto_core gene catalog, then SAMtools (v. 1.16) (Danecek *et al*., 2021) was used to retrieve the number of aligned reads for each sample and the gene length. The resulting aligned reads were normalized in RPKM (reads per kilobase of gene per million mapped reads) for each sample using the following formula:

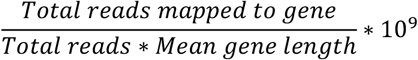

RPKM values were computed for each BGC family in each sample and used to create graphical representations. Only values >0.5 RPKM were selected for further analysis.

A heatmap representing the abundance of BGC families across samples was generated using *heatmap.2* function of the gplots package (v. 3.2.0) (Warners *et al*., 2024) to highlight their distribution patterns. To identify significant BGC families, meaning those differentially abundant between sample groups (CRC/adenoma and healthy), statistical analyses were performed. Specifically, Fisher’s exact test was used to analyze the dispersion of BGC families between groups, and the Wilcoxon rank sum test was applied to compare the distributions of abundance values for all the BGC families between the groups.

Only BGC families with statistically significantly higher RPKM values in the GM of healthy individuals (p<0.05, Wilcoxon rank sum test) were selected for further investigation. The synteny and gene composition of the 4 HM-BGCs were evaluated through detailed analysis of the AntiSMASH (v. 6.1.1) (Blin *et al*., 2021) output files. The organization of adjacent genes was also examined by comparing HM-BGCs with homologous BGCs retrieved from the BGC Atlas (Bağcı *et al*., 2025) and MiBIG (v. 4.0) (Zdouc *et al*., 2025) databases, allowing the reconstruction of the genetic context and the comparison of the organization and putative functions of each cluster. Whenever possible, predictions of BNCs were inferred using PRISM 3 (Skinnider *et al*., 2017) and RiPPMiner (Agrawal *et al*., 2017). The homology between the condensation and AMP-binding domains of HM-BGC3 and the analogous domains of CurF was found by applying SBSPKS v2 (Khater *et al*., 2017).

Two structural models were generated *in silico* using a local installation of AlphaFold2 (AF) (Jumper et al., 2021). The model of the “CurF-like” protein from HM-BGC3 was obtained by providing the full-length sequence (1,513 amino acids) to AF using default parameters. The top-ranking model after relaxation was retained for downstream analyses. Similarly, a structural model of CurF from *L. majuscula* was generated using the protein sequence available in UniProt (accession: Q6DNE7). Firstly, the region of CurF homologous to the “CurF-like” protein was identified via sequence alignment and extracted. This region spans residues 1,990 to 3,154. A CurF model corresponding to this homologous region was then generated by supplying only the selected subsequence to AF, and the highest-confidence relaxed model was again retained.

For the phylogenetic analysis, we used the “CurF-like” protein from HM-BGC3, the CurF protein from the *M. producens* curacin A BGC, and homologous proteins retrieved from the 1,308 BGCs belonging to the BGC Atlas family GCF_14140 (Bağcı et al., 2025). All protein sequences from the BGCs in the GCF_14140 family were extracted together with their corresponding taxonomic annotations and then aligned to the amino acid sequence of the *“CurF-like”* gene from HM-BGC3 using BLASTp (Camacho et al., 2009). Hits showing ≥30% amino acid identity and ≥950 amino acids in length were retained for subsequent analysis. For each BGC, only the best-matching sequence (i.e., the one with the highest identity and length) was kept as representative to avoid redundancy. Filtered sequences were aligned with MAFFT (Katoh & Standley, 2013) using the L-INS-i algorithm, and a maximum-likelihood phylogenetic tree was inferred with IQ-TREE (Minh et al., 2020) under the best-fit substitution model automatically selected by the Bayesian Information Criterion (BIC). Branch support was estimated using 1,000 ultrafast bootstrap replicates. The resulting tree was visualized and annotated in R using ggtree and ggplot2 packages.

Finally, the core genes of the significant BGC families were mapped in a reference population from Rampelli *et al*. (2020) divided as follows: 11 young adults (Y, mean age = 32.2 years), 13 elderly (K, mean age = 72.5 years), 15 centenarians (C, mean age = 100.4 years), and 23 semi-supercentenarians (S, mean age = 106.3 years). In this study, groups C and S were evaluated together as they are representative of individuals with exceptional longevity. The alignment was performed using the same procedure described before, using Bowtie2 (v. 2.3.4.3, with the option ‘--end-to-end’ ‘--very-sensitive’) (Langmead & Salzberg, 2012) to align the metagenome reads of each sample against the constructed proto_core gene catalog and SAMtools (v. 1.16) (Danecek *et al*., 2021) to retrieve the number of aligned reads for each sample and gene length. The number of aligned reads was normalized in RPKM for each sample before graphical representation.

All statistical analyses and graphical representations were performed using R software (v. 4.2.0, www.r-project.org) and the packages gplots (v. 3.2.0) (Warners *et al*., 2024), RcppAlgos (v. 2.9.3) (Wood *et al*., 2025), and xlsx (v. 0.6.5) (Dragulescu & Arendt, 2020). Where not reported, default parameters were used.

## Ancillary Information

### Supporting Information

***Supplementary Table 1. Main features of the metagenome-assembled genomes (MAGs) retrieved from patients with colorectal cancer (CRC), adenoma, and healthy individuals***. For each of the 16,936 MAGs retrieved from 559 fecal samples (258 CRC patients, 81 adenoma patients, and 220 healthy controls), the genome name and size, completeness and contamination percentages, and assigned species are reported. The MAGs from which healthy microbiome (HM)-biosynthetic gene clusters (BGCs) were identified are highlighted in yellow.

**Additional file 1**. Pairwise amino-acid alignment between ctg33_13 (query; HM-BGC3) and CurF.

### Corresponding Author

Further information and request for resources and reagents should be directed to and will be fulfilled by the lead contact, Simone Rampelli.

### Author contributions

**M.C**.: Conceptualization. **D.L**., **D.S**., **S.R**., **C.S**.: Data curation, **D.L**., **D.S**., **S.R**., **C.S**.: Formal analysis, **S.T**., **M.C**., **S.R. P.M**.: Supervision. **D.L**., **S.R**.: Visualization. **D.L**., **M.C**., **S.R**., **S.T**.: Roles/Writing - original draft. **F.D.A**., **T.F**., **D.S**., **S.R**., **P.M**.: Writing - review & editing

## Acknowledgment

This work was supported by the Italian Ministry of University and Research under the National Recovery and Resilience Plan (NRRP), funded by the European Union – NextGenerationEU, through the project “National Biodiversity Future Centre (NBFC)”, and by the PRIN-PNRR 2022 project InBLOOM – INterplay Between Lifestyle and blOod micrObiome in colon rectal cancer prevention: a new Metagenomic perspective.

## Data and code availability

Metagenomic datasets analyzed in this study are publicly available in the repositories cited in the Experimental Section. Accession numbers are provided in the Supporting Information. Any additional data are available from the corresponding author upon reasonable request.

This study does not report original code.

## Abbreviations Used

BGC: biosynthetic gene cluster
BNC: bioactive natural compound
CRC: colorectal cancer
GM: gut microbiome
HGT: horizontal gene transfer
HM-BGC: healthy microbiome-biosynthetic gene cluster
MAG: metagenome-assembled genome
NRPS: non-ribosomal peptide synthetase
PKS: polyketide synthase
RiPP: ribosomally synthesized and post-translationally modified peptide
RRE: RiPP recognizing element.

